# Dual Roles for Nuclear RNAi Argonautes in *C. elegans* Dosage Compensation

**DOI:** 10.1101/2021.07.29.454039

**Authors:** Michael B. Davis, Bahaar Chawla, Eshna Jash, Lillian E. Tushman, Rebecca A. Haines, Györgyi Csankovszki

**Affiliations:** Department of Molecular, Cellular, and Developmental Biology, University of Michigan, Ann Arbor, Michigan, United States of America

**Keywords:** Nuclear RNAi, Argonautes, Dosage Compensation, *C. elegans*, Chromatin structure, Gene regulation, Development, X-linked gene expression, FISH

## Abstract

Dosage compensation involves chromosome-wide gene regulatory mechanisms which impact higher order chromatin structure and are crucial for organismal health. Using a genetic approach, we identified Argonaute genes which promote dosage compensation in *C. elegans*. Dosage compensation in *C. elegans* hermaphrodites is initiated by the silencing of *xol-1* and subsequent activation of the Dosage Compensation Complex (DCC) which binds to both hermaphrodite X chromosomes and reduces transcriptional output by twofold. A hallmark phenotype of dosage compensation mutants is decondensation of the X chromosomes. We characterized this phenotype in Argonaute mutants using X chromosome paint probe and fluorescence microscopy. We found that while nuclear Argonaute mutants *hrde-1* and *nrde-3* exhibit de-repression of *xol-1* transcripts, they also effect X chromosome condensation in a *xol-1*-independent manner. We also characterized the physiological contribution of Argonaute genes to dosage compensation using genetic assays and find that *hrde-1* and *nrde-3*, together with the piRNA Argonaute *prg-1*, contribute to healthy dosage compensation both upstream and downstream of *xol-1*.

## INTRODUCTION

The evolution of sexual dimorphism facilitated the emergence of sex chromosomes in metazoans (Ohno, 1967; Rice, 1984). While the autosomal source of the ancestral sex chromosomes can be divergent between species, a defining factor of sex chromosomes is a difference in the genetic content between the sexes. However, sex chromosomes also bring about the problem of one sex containing half of the gene dose from that chromosome (Ohno, 1967). The problem of haploinsufficiency is evidenced by numerous documented cases of lethality in monosomic conditions (Torres *et al.,* 2008). However, the difference in gene dose associated with sex chromosomes is alleviated by the evolution of dosage compensation (Ohno, 1967). In various organisms, the enormous undertaking of dosage compensation features the regulation of an entire chromosome such that a viable balance of sex chromosome gene expression between the sexes is met. In *C. elegans,* dosage compensation is achieved by a ten-protein complex named the Dosage Compensation Complex (DCC) (For a review, see Lau and Csankovszki, 2015 and Albritton and Ercan, 2018). The DCC is comprised of a pentameric condensin complex, an H4K20me2/3 demethylase (*dpy-21)* and several proteins which confer X chromosome binding specificity (Chuang *et al.,* 1994; Csankovszki *et al.,* 2009; Pferdehirt *et al.,* 2011; Brejc *et al.,* 2017). The DCC downregulates the transcriptional output of both hermaphrodite X chromosomes in interphase cells by two-fold (Kruesi *et al.,* 2013). This system affords a niche opportunity to study the regulatory and functional capacity of condensin in the context of gene regulation.

Indeed, many insights into this unique condensin within the context of the DCC have been discovered. The DCC compacts the X chromosomes, restricting their volume within the nucleus, and facilitates the formation of TADs (topologically associating domains) spaced in ~1Mb intervals along the 17Mb X chromosome (Lau *et al.,* 2014, Crane *et al.,* 2017). The complex is also required for X enrichment of the repressive H4K20me1 mark and loss of the activating H4K16ac mark in interphase (Vielle *et al.,* 2012; Wells *et al.,* 2012; Brejc *et al.,* 2017). Loss of function mutations for the zygotically expressed DCC gene *sdc-2* result in a complete failure for the DCC to load, severe lethality and de-repression of X-linked genes (Dawes *et al.,* 1999). While mutations in the H4K20 histone methyltransferase and demethylase genes lead to significant X chromosome gene de-repression, it is not to the same level as in a loss of DCC function mutation (Kramer *et al.,* 2015; Brejc *et al.,* 2017). An additional pathway facilitates dosage compensation by anchoring the H3K9me2/3 heterochromatin regions of the X chromosome to the nuclear periphery via the nuclear lamina-localized chromo-domain protein CEC-4. Mutants for *cec-4* as well as double mutants for the H3K9me2/3 methyltransferases *met-2 set-25* exhibit modest but significant X-linked gene de-repression (Snyder *et al.,* 2016).

RNAi pathways are a distinct mechanism of controlling gene expression utilizing small interfering RNAs (siRNAs) complexed with Argonaute (AGO) proteins. This RNAi machinery targets mRNAs in the cytoplasm for post-transcriptional silencing or nascent transcripts in the nucleus for co-transcriptional silencing at a gene locus (For a review see Almeida *et al.,* 2019). The AGO family proteins are widely conserved and vary in number across metazoans (Höck and Meister, 2008). As the human genome encodes 8 AGO genes, *Drosophila* have 5, and a fission yeast harbor just one, the striking expansion to 27 AGO genes in *C. elegans* begs the question of their role in gene regulation and relevance to physiology.

The exogenous RNAi pathway in *C. elegans* functions in the silencing of invading viral sequences through the detection of foreign dsRNA, while the endogenous RNAi pathway silences endogenous mRNA transcripts. The piRNA pathway has a conserved role in promoting germline fertility and silencing transposable elements. However the high mismatch pairing capacity for piRNAs and their sheer abundance suggest the piRNA pathway function in *C. elegans* is not fully understood (Lee *et al.,* 2012; Pharad and Theurkauf, 2019). All three branches of the RNAi pathways in *C. elegans* involve several distinct short RNA pools (Ruby *et al,* 2006; Pak and Fire 2007). In the endogenous RNAi silencing pathway, mRNA transcripts for target genes serve as templates for the RNA-dependent RNA Polymerase RRF-3 to generate dsRNA. DICER homolog DCR-1 cleaves this dsRNA into Primary siRNAs which are 26 nucleotides in length with a 5’ bias for guanosine monophosphate (26Gs) (Gent *et al.,* 2009; Han *et al.,* 2009). The primary siRNAs complex with either the ERGO-1(soma and oogenic germline) or ALG-3/4 (spermatogenic gonad) AGOs to target additional mRNA transcripts for the same target gene, upon which many secondary siRNAs are produced (Han *et al.,* 2009; Conine *et al.,* 2010; Vasale *et al.,* 2010). The exogenous RNAi pathway utilizes some of the same machinery, such as DCR-1, which cleaves viral dsRNAs to produce exogenous 26G primary siRNAs. However, exogenously derived primary siRNAs are bound to a distinct AGO called RDE-1, which functions solely in the exogenous RNAi pathway. The primary AGO of the piRNA pathway in *C. elegans* is PRG-1 (Batista *et al.,* 2008; Das *et al.,* 2008). Genomically encoded RNA pol II-transcribed piRNAs are complexed with PRG-1 and target transcripts which are subsequently utilized to generate secondary siRNAs. The exogenous, endogenous and piRNA pathways converge at the synthesis of their respective secondary siRNAs, which complex with a subset of worm-specific Argonautes (WAGOs) to effect post-transcriptional silencing in the cytoplasm, and/or recruit the *nrde* complex to target loci in the nucleus in instances of co-transcriptional silencing (Gent *et al.,* 2010). Secondary siRNAs are 22 nucleotides in length with a 5’ bias for guanosine monophosphate (22Gs) (Ruby *et al,* 2006; Pak and Fire 2007).

The RNAi machinery responsible for co-transcriptional silencing in the nucleus is also required for the accumulation of H3K9me3 and H3K27me3 heterochromatin marks at target loci (Guang *et al.,* 2010; Burkhart *et al.,* 2011; Mao *et al.,* 2015). The corresponding nuclear WAGO genes are *nrde-3* and *hrde-1,* which silence target genes in somatic cells and the germline respectively (Guang *et al.,* 2008; Buckley *et al.,* 2012). The NRDE-3 and HRDE-1 proteins lack the catalytic triad responsible for cleaving mRNAs that is typical of AGO proteins (Yigit *et al.,* 2006). However, both proteins contain an NLS sequence which localizes the WAGO to the nucleus in an siRNA-dependent manner (Guang *et al.,* 2008; Buckley *et al.,* 2012). Mutants for *hrde-1* and *prg-1* show progressive transgenerational sterility at 20 and 25 degrees and are thus designated with the mortal germ line (Mrt) phenotype (Buckley *et al.,* 2012, Simon *et al.,* 2014). Mutants for *prg-1* however, do not exhibit widespread transposable element de-repression. Rather, the transgenerational sterility is likely due to mis-regulation of the downstream WAGOs, which inappropriately re-allocate their silencing capacity to histone mRNAs (Reed *et al.,* 2020).

Here we show that the AGOs *prg-1, hrde-1* and *nrde-3* perform dual roles in *C. elegans* dosage compensation, both in the repression of *xol-1* in hermaphrodites as well as a *xol-1-* independent role. Our FISH experiments indicate that *nrde-*3, *hrde-1,* and *prg-1* mutants exhibit an X chromosome decondensation phenotype and our XO rescue experiments indicate that these genes contribute physiologically to dosage compensation. We demonstrate that while both the AGO-mediated XO rescue and X chromosome compaction are *xol-1-*independent, the effects of these mutations on hermaphrodite viability are largely *xol-1*-dependent. We present a model whereby *nrde-3* and *hrde-1* facilitate the transcriptional repression of *xol-1* possibly as the effectors of *prg-1-*21UX1 piRNA pathway, and downstream of *xol-1* these nuclear AGOs promote the compaction of X chromosomes in a separate but physiologically relevant role.

## MATERIALS AND METHODS

### *C. elegans* strains and maintenance

Strains were grown under standard conditions as described (Brenner, 1974). Worm strains include: N2; SX922 *prg-1(n4357)* I; EKM 200 *prg-1(n4357)* I *lon-2(e678) xol-1(y70)* X; WM157 *wago-11(tm1127)* II; YY470 *dcr-1(mg375)* III; EKM 89 *hrde-1(tm1200)* III; EKM 101 *him-8(e1489)* III *mIs11* IV; EKM 152 *hrde-1(tm1200)* III *lon-2(e678) xol-1(y70)* X; EKM 117 *hrde-1(tm1200)* III *mIs11* IV; EKM 177 *mIs11* IV *nrde-3(tm1116)* X; EKM 178 *nrde-3(tm1116) lon-2(e678) xol-1(y70)* X; EKM 176 *mIs11* IV *him-5(21490)* V *nrde-3(tm1116)* X; WM191 MAGO12 *sag-2(tm894) ppw-1(tm914) wago-2(tm2686) wago-1(tm1414)* I, *wago-11(tm1127) wago-5(tm1113) wago-4(tm1019)* II, *hrde-1(tm1200) sago-1(tm1195)* III, *wago-10(tm1186)* V, *nrde-3(tm1116)* X; CB1489 *him-8(e1489)*IV; TY4403 *him-8(e1489) xol-1(y9) sex-1(y263)*; WM158 *ergo-1(tm1860)* V; WM27 *rde-1(ne219)* V; WM156 *nrde-3(tm1116)* X; EKM 125 *lon-2(e678) xol-1(y70)* X. Worms were fed OP50. In all FISH experiments (Figure 1 and 4), Hermaphrodite viability (Figure 5) and XO rescue (Figure 6) experiments, worms for every genotype were grown at 15°C for all experiments. This measure was taken to control for temperature-sensitive effects in strains containing *hrde-1* and *prg-1* mutations.

**Figure 1.**
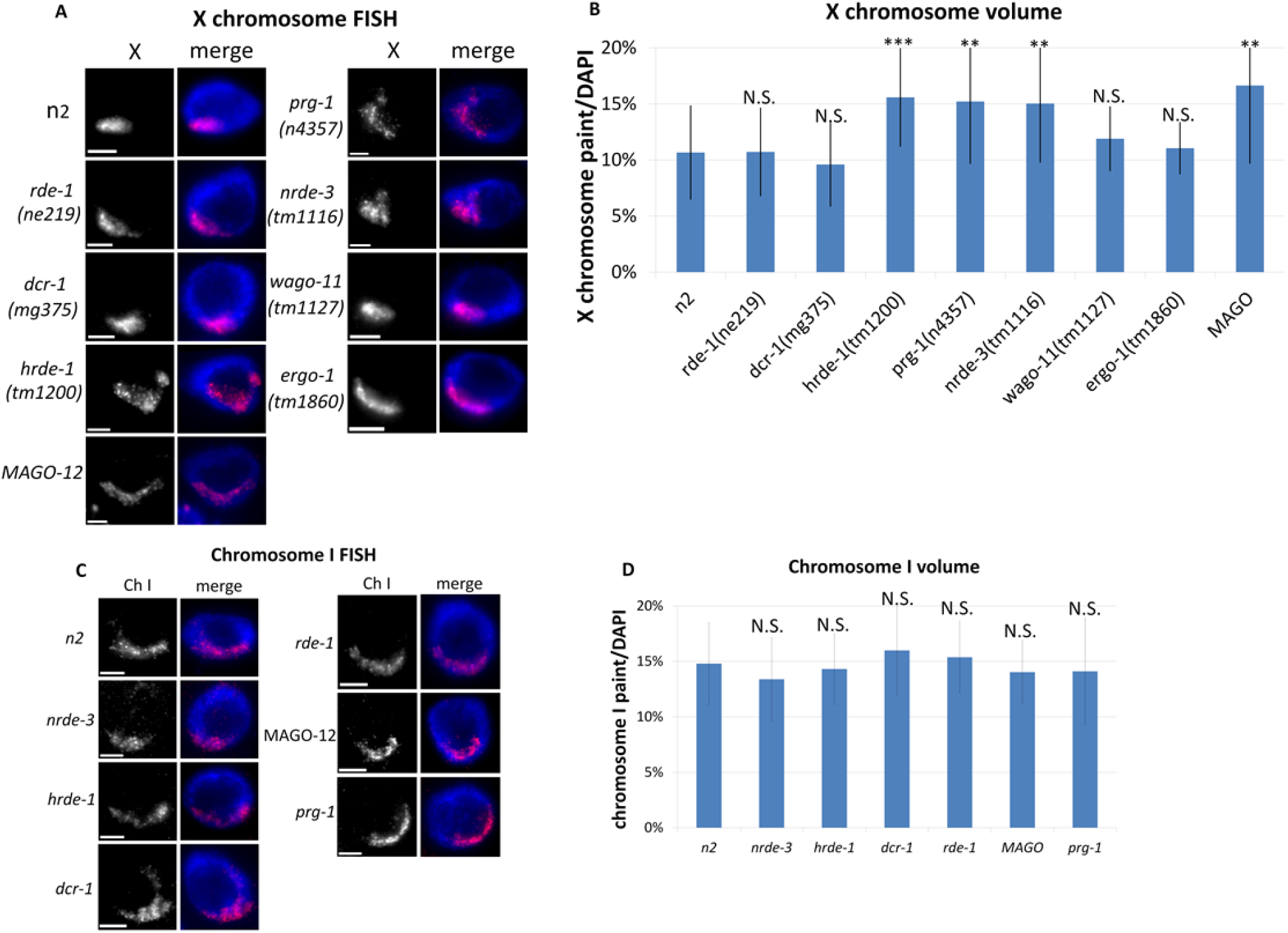
X chromosome is de-condensed in nuclear RNAi Argonaute mutants. **(A)** FISH X chromosome paint (red) with nuclear (DAPI) staining is shown in adult hermaphrodite intestinal cells. The X paint signal region is larger in *nrde-3, hrde-1, prg-1*, and *MAGO-12* mutants. **(B)** Quantification for X chromosome volume: (X chromosome paint voxels/DAPI voxels). **(C)** FISH chromosome I paint (red) with nuclear (DAPI) staining is shown. The Ch I paint signal region in wild type (N2) and each mutant with an X chromosome phenotype is similar for this autosome control. **(D)** Quantification for chromosome I volume: (chromosome I paint voxels/DAPI voxels). Significance for Band D is based on a Student’s T Test: **=p<0.01, ***= p<0.001 (N2 compared to each mutant). Error bars represent SD.

### Male rescue and hermaphrodite viability assays

#### RNAi-based male rescue assay

Worms with the genotype *him-8(e1489); xol-1(y9) sex-1(y263)* were fed *E. coli* strain HT115 expressing double stranded RNAi for each gene of interest. Bacteria were picked from a single colony and grown overnight for 8-10 hours at 37°C. 150ul of bacteria were seeded on NMG plates with IPTG (0.2%w/v) and Carbenicillin (1ug/ml) and worms were transferred to plates after one day. The male rescue assay in Figure 2 was conducted as outlined in (Petty *et al.,* 2009). *him-8(e1489) xol-1(y9) sex-1(y263)* L1 hatchlings were transferred to RNAi plates and allowed to grow until L4 before (2-3) worms were transferred to new RNAi plates. Worms were allowed to lay eggs for 24 hours at 20°C before parents (P0) were removed. Embryos were counted and the presence of males (in the F1) was scored after 2-4 days of growth at 20°C. As a hermaphrodite with the *him-8(e1489)* mutation produces a population of 38% males, the % of males rescued was calculated by dividing the number of males observed by the number of males expected (expected males = total eggs laid X 0.38) (Philips *et al.,* 2005). Statistical significance was evaluated using Chi square tests for each comparison of two conditions. The null hypothesis was that no significant difference in male rescue would be found between each condition measured. The expected ‘viable male worms’ was calculated by taking the summed % viable male worms between the two groups being tested and multiplying that proportion by the total male worm number for each corresponding condition.

#### XO rescue assay

For the XO rescue assays on Figure 6B, crosses were set up between *lon-2 xol-1* hermaphrodites with the appropriate additional mutation (*nrde-3* or *hrde-1*) and *mIs11* males with the appropriate additional mutation. F1 cross progeny were identified by the presence of the GFP transgene. XX cross progeny were *lon-2*/+, and therefore developed as non-Lon hermaphrodites. XO cross progeny were hemizygous for *lon-2*, and developed as Lon males or hermaphrodites. F1 progeny were scored for the presence of GFP+ males and/or GFP+ Lon hermaphrodites (XO cross-progeny), as well as the presence of GFP+ non-Lon hermaphrodites (XX cross-progeny). The % of XO rescued animals was calculated by dividing the number of rescued XO animals (GFP+ males and GFP+ Lon hermaphrodites) observed by the number of XX (GFP+ non-Lon) hermaphrodites. This calculation assumes that equal numbers of XX and XO cross-progeny are produced from a mating of a male to a hermaphrodite.

For the XO rescue on empty vector (EV) or *sex-1(RNAi), lon-2 xol-1* mutants with the appropriate additional mutation (*nrde-3* or *hrde-1*) were moved to *sex-1(RNAi)* or (EV) plates as hatchlings. Simultaneously, *mIs11* worms with the appropriate additional mutation (*nrde-3* or *hrde-1*) were also moved to *sex-1(RNAi)* plates as hatchlings. After approximately two days of growth at 15°C, L4’s were selected from both genotypes to initiate crosses on new *sex-1(RNAi)* or *EV* plates. F1 progeny were scored as above. Statistical significance was evaluated using Chi square tests for each comparison of two conditions. The null hypothesis was that no significant difference in XO rescue would be found between each condition measured. The expected ‘viable XO worms’ calculation was formulated by taking the summed % viable XO worms between the two groups being tested and multiplying that proportion by the total XO worm number for each corresponding condition.

#### Hermaphrodite viability assay

For the hermaphrodite viability assays, strains of the appropriate genotype were moved to *sex-1(RNAi)* or *EV* plates at the L1 stage. After two days of growth at 15°C L4’s were picked and transferred to new RNAi plates and allowed to lay embryos for 24 hours before being transferred again to new plates. Following the removal of parents each day the number of embryos on each plate was counted and after two full days at 15°C viability was scored based on the (number of live worms) divided by the (number of eggs laid). Statistical significance was evaluated using Chi square tests for each comparison of two conditions. The null hypothesis was that no significant difference in hermaphrodite viability would be found between each condition measured. The expected ‘alive’ calculation was formulated by taking the summed % viability between the two groups being tested and multiplying that proportion by the sample size (*n*) for each corresponding condition.

### Fluorescence In Situ Hybridization (FISH)

FISH probe DNA templates were created through degenerate oligonucleotide-primed PCR reactions to amplify purified yeast artificial chromosome (YAC) DNA corresponding to either the X chromosome or chromosome I (Csankovszki *et al.,* 2004, Nabeshima *et al.,* 2011). dCTP-Cy3 (GE) was incorporated along with standards dNTPs for visualization purposes using random priming (Labeling 5X Buffer Promega). For the fixation, worms were dissected in 1X sperm salts (50 mM Pipes pH 7, 25 mM KCl, 1 mM MgSO4, 45 mM NaCl and 2 mM CaCl2) and fixed in 1.6% PFA on slides for 5 minutes at RT. Subsequently slides were frozen on dry ice to 20 minutes before a series of 3X 10 minute washes in PBST. Slides were then washed in an ethanol series: 70%, 80%, 95%, 100% for 2 minutes each. After allowing slides to dry for 5 minutes, 10ul of Cy-3-labelled probe was applied with a coverslip and heated to 95°C for 3 minutes. Slides were then incubated in a humidity chamber overnight at 37°C. The following day slides were washed in 2X SSC + 50% formamide (3X 5-minute washes); 2X SSC (3X 5 minute washes); 1X SSC (10 minutes); 4X SSC+DAPI (10 minutes). Slides were then mounted with Vectashield (Vector Labs).

### Imaging and Quantification

Images were taken with a Hamamatsu ORCA-ER digital camera mounted on an Olympus BX61 epi-fluorescence microscope with a motorized Z drive. The 60X APO oil immersion objective was used for all images. Series of Z stack images were collected (stack size 0.2um) and all images shown are projection images summed from ~3 microns. Quantification was conducted in the Slidebook 5 program (Intelligent Imaging Innovations). Using the mask function, segment masks were drawn individually for the DAPI signal and Cy3 signal for each nucleus. Within the statistics menu the mask statistics function as selected with DAPI as the primary mask and Cy3 as the secondary mask. The morphometrix (voxels) and crossmask (voxels) were selected for the analysis. These procedures generated a number for the overlap corresponding to the volume of the Cy3 and DAPI masks from which chromosome volume was calculated using Cy3 volume/DAPI volume. The average volume of Cy3/DAPI for all worms of a given genotype was calculated and an unpaired (2-sample) Student’s T-test was performed to compare the means of each genotype to the appropriate control. At least 20 nuclei per condition were quantified and identical probe batches were used for each experiment with a wild type (N2) control.

### qRT-PCR

Synchronous culture of gravid adult worms was grown at 25°C to induce temperature-dependent gene expression changes in the temperature sensitive mutants. Worms were then bleached to obtain embryos. Embryos were lysed using a bead beater with 0.1mm zirconia/silica beads (BioSpec Products cat. no. 11079101z). TRIzol-chloroform (Invitrogen, Fisher Scientific) separation of the samples was followed by RNA extraction using the QIAGEN RNeasy Mini Kit with on-column DNase I digestion. cDNA was generated from extracted RNA using random hexamers with SuperScript III Reverse Transcriptase (Invitrogen). RT-qPCR reaction mix was prepared using Power SYBR Green Master Mix (Applied Biosystems) with 10μl SYBR master mix, 0.8μl of 10μM primer mix, 2μl sample cDNA, and 7.2μl H2O. Samples were run on the Bio-Rad CFX Connect Real-Time System. Log2 fold change was calculated relative to control (*cdc-42*). Control primers were taken from Hoogewijs *et al.,* (2008). Statistical significance was calculated using two-tailed unpaired Student’s t-test. Error bars represent standard deviation.

Primer sequences: *xol-1* forward GGTCCCACCAGAAATGCAG, *xol-1* reverse – CTGTTTCCATGTAAGTTAAGACTGG, *cdc-42* forward – ctgctggacaggaagattacg, *cdc-42* reverse – ctcggacattctcgaatgaag.

### Immunofluorescence

Immunofluorescence experiments were performed as described (Csankovszki *et al.,* 2009). One day post-L4 worms were dissected in 1X sperm salts (50 mM Pipes pH 7, 25 mM KCl, 1 mM MgSO4, 45 mM NaCl and 2 mM CaCl2) and fixed in 2% paraformaldehyde in 1X sperm salts for 5 minutes with a coverslip. Slides were frozen on dry ice for at least 15 minutes. Coverslips were removed and slides were washed in PBS with 0.1% Triton X-100 (PBST) (3X 10 minutes each) and subsequently incubated with primary rabbit anti-H3K9me3 (Active Motif #39765) antibodies (at 1:500 dilution) with a square parafilm. Slides were then moved to a humidity chamber overnight at room temperature. Slides were then washed in PBST (3X 10 minutes each) and secondary donkey anti-rabbit-FITC (Jackson Immunoresearch) antibody at 1:100 dilution was added with a square parafilm slip. Slides were incubated for 2 hours at 37°C before another series of washes in PBST (3X 10 minutes each). The final PBST wash included DAPI. Slides were mounted with Vectashield (Vector Labs). Significance for H3Kme3 signal pattern differences between genotypes was calculated by Chi square tests conducted between two conditions for each comparison under the null hypothesis of no significant difference in H3K9me3 pattern between genotypes.

## RESULTS

### X chromosome decondensation in RNAi mutants

A hallmark phenotype of mutants for DCC genes and additional genes influencing dosage compensation is a decondensed X chromosome territory (Lau *et al.,* 2014; Snyder *et al.,* 2016; Brejc *et al.,* 2017). We previously found that heterochromatin proteins, including the H3K9 methyltransferases MET-2 and SET-25, and the chromodomain-containing protein CEC-4 that bind H3K9me, promote dosage compensation and X chromosome compaction (Snyder *et al.,* 2016). Since nuclear RNAi can lead to deposition of H3K9me3 and heterochromatin formation, we wondered whether nuclear RNAi genes would have similar effects. In a previous study, we reported that in nuclear RNAi mutants *morc-1* and *hrde-1* the X chromosomes were decondensed (Weiser *et al.,* 2017). To determine which RNAi pathways influence X chromosome compaction, we conducted X chromosome paint fluorescence in situ hybridization (FISH) experiments with loss of function mutants for various genes in the *C. elegans* RNAi pathways (Figure 1A). Our analysis included mutants for genes eliminating function specifically in each branch of the RNAi pathways (exogenous, endogenous and piRNA) as well as mutants for downstream AGO genes which are shared by multiple pathways. The exogenous RNAi pathway was represented by *rde-1(ne219)*, a mutation for the primary AGO which disrupts silencing in the pathway (Tabara *et al.,* 1999). Endogenous RNAi pathway mutations included: a mutation in DICER homolog *dcr-1(mg375),* a helicase domain mutant impairing the production of primary 26G siRNAs specifically in the endogenous RNAi pathway, as well as *ergo-1(tm1860)*, a null mutant for the primary AGO of the endogenous RNAi pathway which eliminates 26G primary siRNAs (Bernstein *et al.,* 2001; Yigit *et al.,* 2006; Pavelec *et al.,* 2009; Welker *et al.,* 2010). The piRNA pathway mutation included *prg-1(n4357)*, a null mutant for the primary AGO of the pathway (Batista *et al,* 2008; Das *et al,* 2008). Downstream AGO mutations included *hrde-1(tm1200), nrde-3(tm1116)* and *wago-11(tm1127)*, which represent null mutants for three of the four nuclear AGO encoding genes, as well as the MAGO-12 (Multiple AGO) mutant bearing loss of function mutations in all 12 of the WAGO genes (*C. elegans* deletion consortium *et al.,* 2002; Guang *et al,* 2008; Buckley *et al,* 2012; Gu *et al.,* 2009).

In intestinal cells of wild type hermaphrodites with intact dosage compensation, the X chromosomes occupied about 10% of the nuclear volume on average, consistent with previous studies (Figure 1B) (Lau *et al.,* 2014; Snyder *et al.,* 2016; Brejc *et al.,* 2017). The *hrde-1, nrde-3, prg-1,* and MAGO-12 mutants displayed significantly decondensed X chromosome volumes, occupying about 15-16% of nuclear volume, and in the range of what was previously reported for heterochromatin mutants (Figure 1B) (Snyder *et al.,* 2016). The X chromosome decondensation phenotype was absent from the *rde-1, dcr-1,* and *ergo-1* mutants, which suggests the effect is not mediated through the exogenous or the endogenous RNAi pathway. The less-characterized nuclear AGO mutant *wago-11* was also not significantly different from wild type. While the *hrde-1, nrde-3, prg-1,* and MAGO-12 mutants exhibit significantly increased X chromosome volume compared to wild type animals, they were not significantly different in X chromosome volume from each other (Figure 1B). Moreover, the MAGO-12 mutant exhibited the same degree of X chromosome decondensation as the single AGO mutants. Based on these results, we hypothesize that the contribution from these mutants to X chromosome compaction is likely a common pathway involving a subset of the RNAi machinery.

To identify whether the X chromosome decondensation observed in the nuclear RNAi AGO mutants was specific to the X chromosome, we also conducted FISH experiments with whole chromosome I paints in each mutant (Figure 1C). Each of the RNAi mutants displayed chromosome I volumes similar to wild type (Figure 1D). Chromosome I occupied about 15% of the nuclear volume in wild type N2, and the volume in mutants ranged from 13.4% in *nrde-3* to about 16% in *dcr-1* (Figure 1D). The variation observed in each mutant was not significantly different from wild type worms, highlighting that the X chromosome volume decondensation phenotype is not a genome-wide feature of the nuclear RNAi AGO mutants.

### Depletion of nuclear RNAi AGOs rescues ***xol-1*** mutant males

To further define the individual contribution of AGO mutations to dosage compensation we conducted male rescue experiments. The assay is based on the principle that while dosage compensation is required for XX hermaphrodite viability, if ectopically activated in XO males, dosage compensation is lethal (Miller *et al.,* 1988, Petty *et al.,* 2012). Mutation in the *xol-1* master switch gene bypasses the innate chromosome counting mechanism in *C. elegans,* effectively activating dosage compensation in a sex-independent manner (Rhind *et al.,* 1995). *xol-1* males are 100% lethal due to the inappropriate activation of dosage compensation acting on their sole X chromosome. However, *xol-1* mutant males can be rescued by inactivation of a gene required for dosage compensation (see methods for details). We previously reported that depletion nuclear RNAi genes *morc-1* and *hrde-1* led to small, but significant rescue of *xol-1* males (Weiser *et al.,* 2017). Here we report that in addition to *hrde-1,* RNAi of *prg-1* and *nrde-3* also rescued males with ectopic dosage compensation (Figure 2A and B). On empty vector only about 1% of males are rescued, while a positive control knockdown of DCC component *dpy-27* yielded a rescue of 52% of males. *prg-1(RNAi)* rescued 17.5% of males, *hrde-1(RNAi)* rescued 18.8% of males, and *nrde-3(RNAi)* rescued 6.1% of males (Figure 2A and B). Thus, the nuclear RNAi AGOs and the piRNA AGO *prg-1* promote X chromosome dosage compensation.

**Figure 2.**
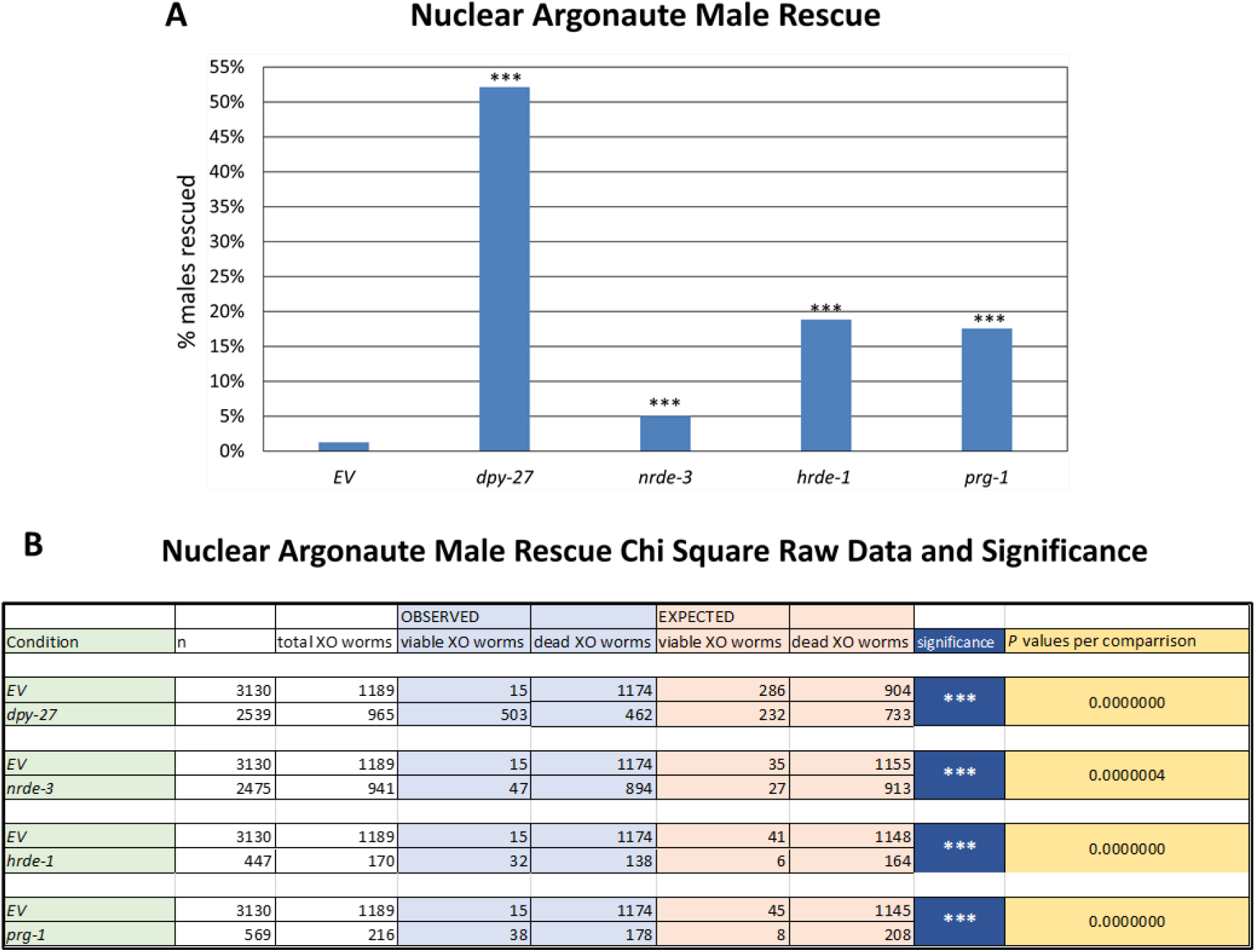
RNAi screen implicates nuclear Argonautes in dosage compensation. **(A)** RNAi knockdown for various genes in a *him-8; xol-l sex-1* background. Percent of rescued males is indicated. DCC component gene *dpy-27* rescues a large % of males and the other genes shown rescue a smaller but reproducible number of males. Significance is based on Chi square analysis: *= p<0.05, **=p<0.01, ***= p<0.001. (B) Raw data from the Chi Square analysis.

### ***xol-1*** is de-repressed in ***hrde-1*** and ***nrde-3*** mutants

An earlier study from Tang *et al.,* (2018) reported a role for *prg-1* in repression of *xol-1.* The study identified a piRNA *21-ux1* which, when complexed with *prg-1*, is partially responsible for repressing *xol-1* in hermaphrodites to permit dosage compensation to turn on. They showed that both *xol-1* transcripts and protein levels are de-repressed in hermaphrodites mutant for *prg-1* as well as mutants with the *21-ux1* piRNA deleted. We conducted RT-qPCR on *hrde-1* and *nrde-3* and *prg-1* mutants to determine whether *xol-1* was similarly de-repressed in the nuclear RNAi AGO mutants (Figure 3). *hrde-1* and *nrde-3* mutants both exhibited significant *xol-1* derepression (Figure 3). The level of derepression in *nrde-3* mutants was similar to *prg-1* mutants, and somewhat less in *hrde-1* mutants. As the canonical piRNA pathway involves nuclear RNAi AGO-mediated silencing, this result suggests that repression of *xol-1* by piRNAs may be reinforced by nuclear RNAi (Ashe *et al.,* 2012; Bagijn *et al.,* 2012; Lee *et al.,* 2012; Shirayama *et al.,* 2012). Thus *nrde-3, hrde-1,* and *prg-1* contribute to dosage compensation upstream of *xol-1* through its repression. Intriguingly, however, our male rescue assay was conducted in a *xol-1* null mutant background, thus the male rescue data (Figure 2) suggests that these genes may play an additional *xol-1-*independent role to promote dosage compensation.

**Figure 3.**
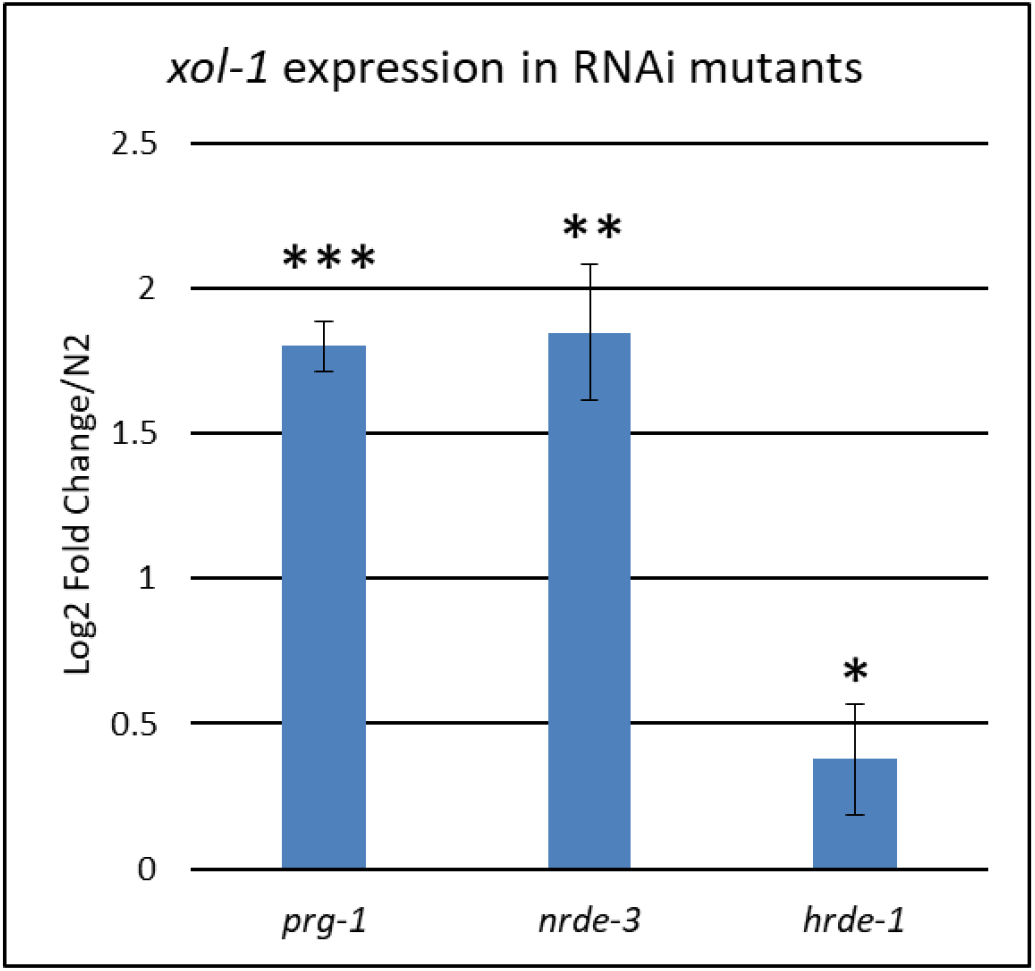
*Xol-1* is de-repressed in *hrde-1* and *nrde-3* mutants. Bar graph shows qRT-PCR results. *xol-1* mRNA transcript levels normalized to wild type (N2). *prg-1* is shown as a positive control. *xol-1* transcripts are significantly more abundant in *hrde-1* and *nrde-3. xol-1* is de-repressed to a similar level in *prg-1* and *nrde-3* mutants, and to a lesser extent in *hrde-1.* Significance is based on a Student’s T-Test: **=p<0.01, ***= p<0.001. Error bars represent SD.

### X chromosome decondensation in nuclear RNAi AGO mutants is independent of ***xol-1***

Since *prg-1, hrde-1 and nrde-3* contribute to the repression of *xol-1* (Figure 3), and our male rescue data (Figure 2) show that RNAi of these genes also rescues males in a *xol-1-* independent manner, the X chromosome compaction defect in the mutants (Figure 1) may or may not be *xol-1-*dependent. To determine if the X chromosome decondensation in *hrde-1, nrde-3,* and *prg-1* mutants (Figure 1) is due to *xol-1* de-repression or a *xol-1-*independent role, we conducted X chromosome FISH in double mutants for *xol-1* and each of the AGO genes (Figure 4A). While *xol-1* mutants did not exhibit X chromosome decondensation, in each double mutant condition, the X chromosome decondensation phenotype of *hrde-1, nrde-3* and *prg-1* persisted. These results suggest that the *xol-1* repression role of the AGO genes is separate from the X chromosome compaction role (Figure 4 B). We also assessed the volume of Chromosome I in each of the nuclear AGO double mutants with *xol-1* and found no significant difference from wild type animals or *xol-1* mutants (Figure 4 C and D). Thus the chromosome decondensation phenotype is X-specific and *xol-1*-independent.

**Figure 4.**
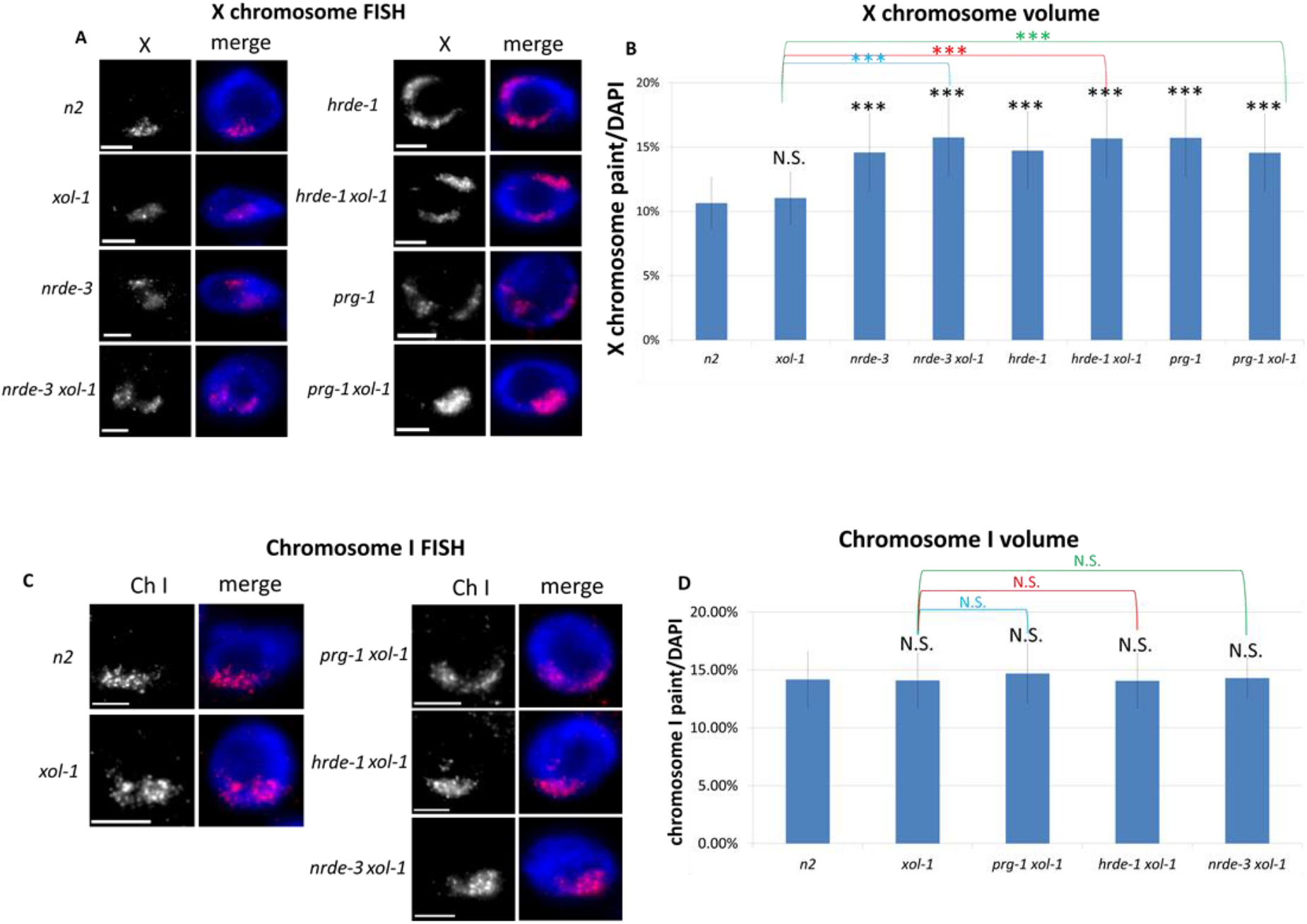
nuclear RNAi Argonaute mutant X de-condensation is independent of *xol-1.* **(A)**FISH X chromosome paint (red) with nuclear (DAPI) staining is shown in adult hermaphrodite intestinal cells. The X paint signal region in single mutants for each Argonaute(*nrde-3, hrde-1, prg-I*) is larger than wild type (N2) and *xol-1*, and similar to its double mutant counterpart with *xol-1*. **(B)** Quantification for X chromosome volume: (X chromosome paint voxels/DAPI voxels). **(C)** FISH chromosome I paint (red) with nuclear (DAPI) staining is shown. The Ch I paint signal region in wild type (N2) and *xol-1* negative controls are similar to each Argonaute mutant with *xol-1*. **(D)** Quantification for chromosome I volume: (chromosome I paint voxels/DAPI voxels). Significance for B and D is based on a Student’s T Test: **=p<0.01, ***= p<0.001 (Black asterisks and labels are N2 compared to each mutant; colored asterisks and labels are *xol-1* compared to each double mutant). Error bars represent SD.

### Synergistic lethality of nuclear RNAi AGO mutants with ***sex-1(RNAi)***

The dual role for nuclear RNAi AGO genes is akin to that of *sex-1,* a better characterized gene in dosage compensation. *sex-1* is an X-chromosome encoded “X signal element” (XSE) which negatively regulates *xol-1* expression, while also contributing to dosage compensation downstream of *xol-1* through an uncharacterized mechanism (Gladden *et al.,* 2007). Hermaphrodite viability data indicate a high degree of lethality in *sex-1* null mutants (~20-37% viable) (Carmi *et al.,* 1998, Gladden *et al.,* 2007). This lethality is largely due to *xol-1* derepression, as x*ol-1 sex-1* double mutants are much more viable (~85% viability) (Gladden *et al.,* 2007). However, a mutation in *xol-1* does not completely rescue lethality, indicating that *sex-1* also plays a *xol-1*-independent role promoting hermaphrodite viability (Gladden *et al.,* 2007). In order to determine the extent to which *hrde-1* and *nrde-3* contribute to hermaphrodite viability upstream and downstream of *xol-1*, we conducted hermaphrodite viability assays with and without the *xol-1(y9)* mutation and *sex-1(RNAi)* (Figure 5). Briefly, worms were subjected to *sex-1(RNAi)* from the L1 larval stage. For each experiment, we counted the total number of eggs laid, the number of unhatched (dead) eggs, the number of dead worms, and the number of live worms that were alive two days after the eggs were laid. The proportion of live worms is shown on Figure 5.

*nrde-3* and *hrde-1* hermaphrodite viability was similar to wild type (N2) on control empty vector RNAi, although the viability of *nrde-3* mutants was somewhat reduced (Figure 5A). However on *sex-1(RNAi),* both mutants exhibited enhanced lethality (Figure 5B, Table S1). Enhanced lethality in the *sex-1* mutant background is a characteristic of genes promoting dosage compensation (Gladden *et al.,* 2007). Thus, this data is consistent with a role for *hrde-1* and *nrde-3* in dosage compensation. Much of the discrepancy between hermaphrodite viability in wild type N2 versus *nrde-3* and *hrde-1* was eliminated with the addition of *xol-1* mutation to each of the single mutants, suggesting that regulation of *xol-1* is an important component of this effect. However, there was still a small, but significant increase in lethality on *sex-1(RNAi)* for *hrde-1 xol-1* and *nrde-3 xol-1* compared to *xol-1* (Figure 5B, Table S1). These results indicate that the nuclear AGO genes contribute to hermaphrodite viability similar to *sex-1*, to a greater extent upstream of *xol-1,* and to a lesser but significant extent downstream of *xol-1.*

**Figure 5.**
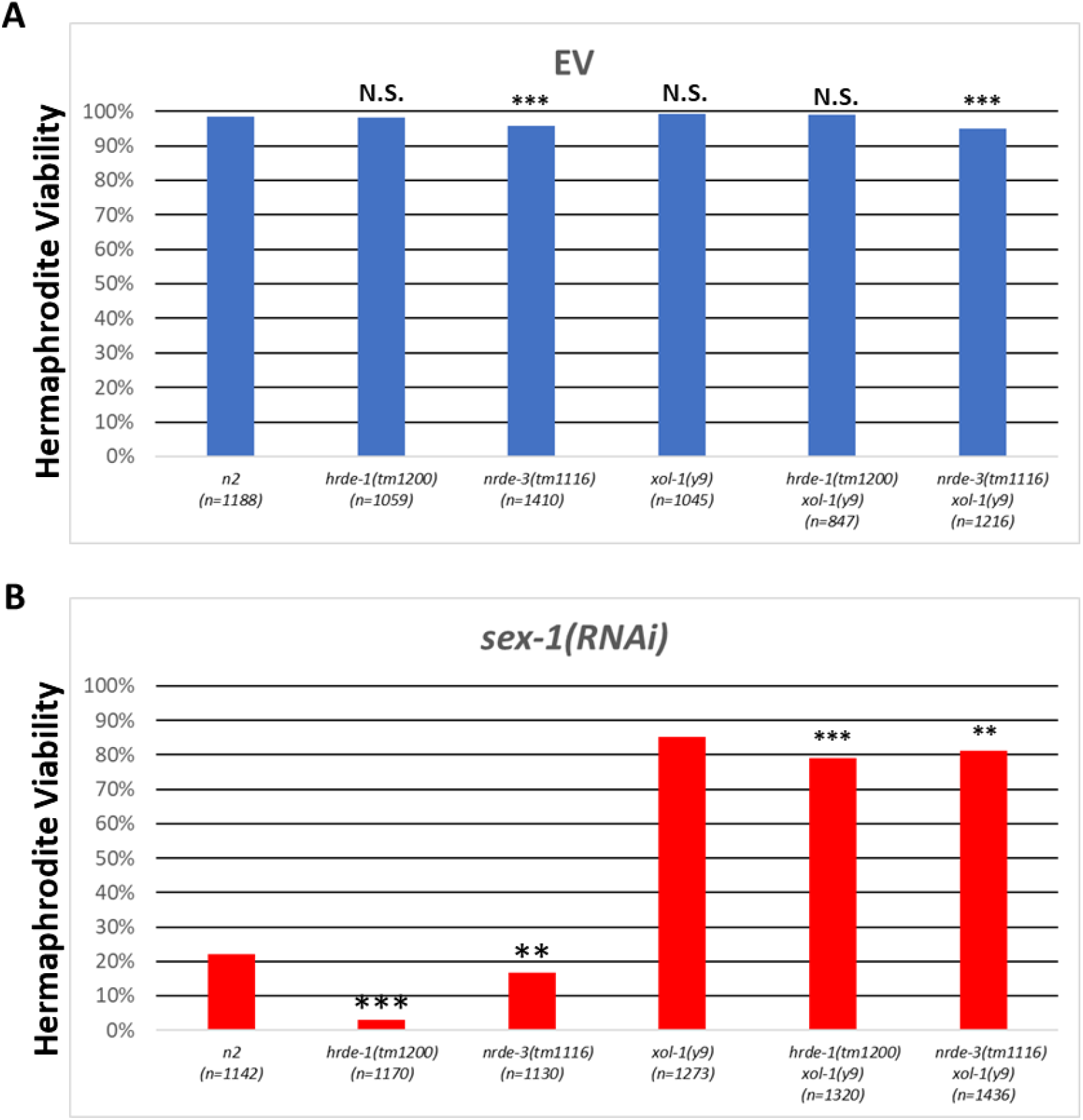
Hermaphrodite Viability of nuclear Argonaute mutants. **(A)** Hermaphrodite viability on empty vector RNAi. % viability (calculated as the number of live worms/total number of eggs laid) is shown. **(B)** Hermaphrodite viability of *hrde-1* and *nrde-3* mutants with and without *xol-1* on *sex-1(RNAi). hrde-1* mutants are only 3% viable and *nrde-3* mutants are slightly less viable than wild type (N2) worms. The lethality is *xol-1* dependent. Significance in A and B are based on comparisons of N2 to each single mutant *(hrde-1* or *nrde-3*) and *xol-1* to each double mutant. Significance is based on *p* Values from Chi Square analysis for each comparison *=p<0.05, **=p<0.01, ***= p<0.001 (See supplemental table 1).

### Nuclear RNAi AGO mutant XO rescue with ***sex-1(RNAi)***

While hermaphrodite viability defects are not a direct readout of dosage compensation, the rescue of *xol-1* mutant XO animals is a highly specific readout of dosage compensation function. To better understand the physiological contribution of *hrde-1* and *nrde-3* downstream of *xol-1,* we refined our XO rescue assay, adopting an approach modified from Gladden *et al.,* (2007) which utilizes a marker mutation to directly score for the XO karyotype, rather than the male phenotype (Figure 6A). In our RNAi-based *xol-1* male rescue assay (Figure 2) we scored the percentage of rescued worms based on the male phenotype. However, because the function of XOL-1 is also needed for male development, there could be candidate genes which yield rescued XO worms with the hermaphrodite phenotype. The strain used for the RNAi assay also has a mutation in *sex-1* which partially disrupts dosage compensation and sensitizes the strain for rescue. Our modified *xol-1* male rescue assay takes advantage of the recessive X-linked *lon-2(e678)* mutation, which renders hemizygous XO or homozygous XX mutant worms ~50% longer than wild type or *lon-2*/+ XX animals. Thus, with the *lon-2(e678)* marker we eliminate these potential false negatives by directly scoring the karyotype. The assay relies on a mating between a male and hermaphrodite which generates a theoretical yield of 50% males to 50% hermaphrodites (Figure 6 A). To distinguish cross-progeny from self-progeny, we include an autosomal linked GFP marker (*mIs11)* in the males of the cross, resulting in GFP+ cross progeny and GFP- self-progeny. The assay also assesses the rescue impact of loss of function mutants, rather than RNAi-based variable knockdown for candidate genes (Figure 6 A). The genetic background of the control cross is thus *lon-2(e678) xol-1(y9)* hermaphrodites crossed to *mIs11* males, whereby *hrde-1,* and *nrde-3* mutations were separately crossed into both of the parent strains for each experiment to ensure that the parent and progeny are homozygous mutant for the gene assessed (for details see Methods). For control, we analyzed the effects of mutations in the DCC subunit and H4K20 demethylase *dpy-21*. Mutations in *dpy-21* rescue a large proportion of XO animals. *hrde-1* and *nrde-3* mutations independently did not significantly rescue XO worms -*ctrl (0.00%)*; *hrde-1 (0.23%); nrde-3 (1.05%)* (Figure 6 B, Table S2). Since reduced *sex-1* function enhances rescue of *xol-1* males (Figure 2), we repeated the rescue matings on *sex-1(RNAi).* On *sex-1(RNAi)* we observed a significant degree of rescue in *hrde-1* and *nrde-3* mutants compared to *sex-1(RNAi)* alone: *sex-1(RNAi) (0.00%); nrde-3 (16.80%), hrde-1 (9.35%)* (Figure 6 C). Thus, while *nrde-3* and *hrde-1* mutations alone are insufficient to rescue XO progeny to a developmental stage that can be scored in this assay, in the sensitized background of *sex-1(RNAi)* their role becomes evident. The overwhelming majority of rescued animals were Lon male, with the Lon hermaphrodite phenotype occurring in four or fewer worms in each condition. Moreover, Lon hermaphrodites were not observed at all in the *nrde-3* and *hrde-1 sex-1(RNAi)* conditions possibly due to *sex-1’s* role sex determination (Gladden *et al.,* 2007). Interestingly, the XO rescue of *hrde-1* and *nrde-3* worms slightly increased on control empty vector alone compared to standard non-RNAi food (Figure 6B and C). One possibility is that engagement of the RNAi machinery with empty vector in the background of nuclear AGO mutations disrupts dosage compensation to a small degree. Overall, these results suggests that in addition to regulating *xol-1*, *nrde-3* and *hrde-1* also promote dosage compensation downstream of *xol-1*. We were unable to assess the role of *prg-1* in this assay, due to the severely reduced fertility of *prg-1* mutant males. However, the RNAi-based rescue (Figure 2) indicates that *prg-1* likely also plays a dual role.

**Figure 6.**
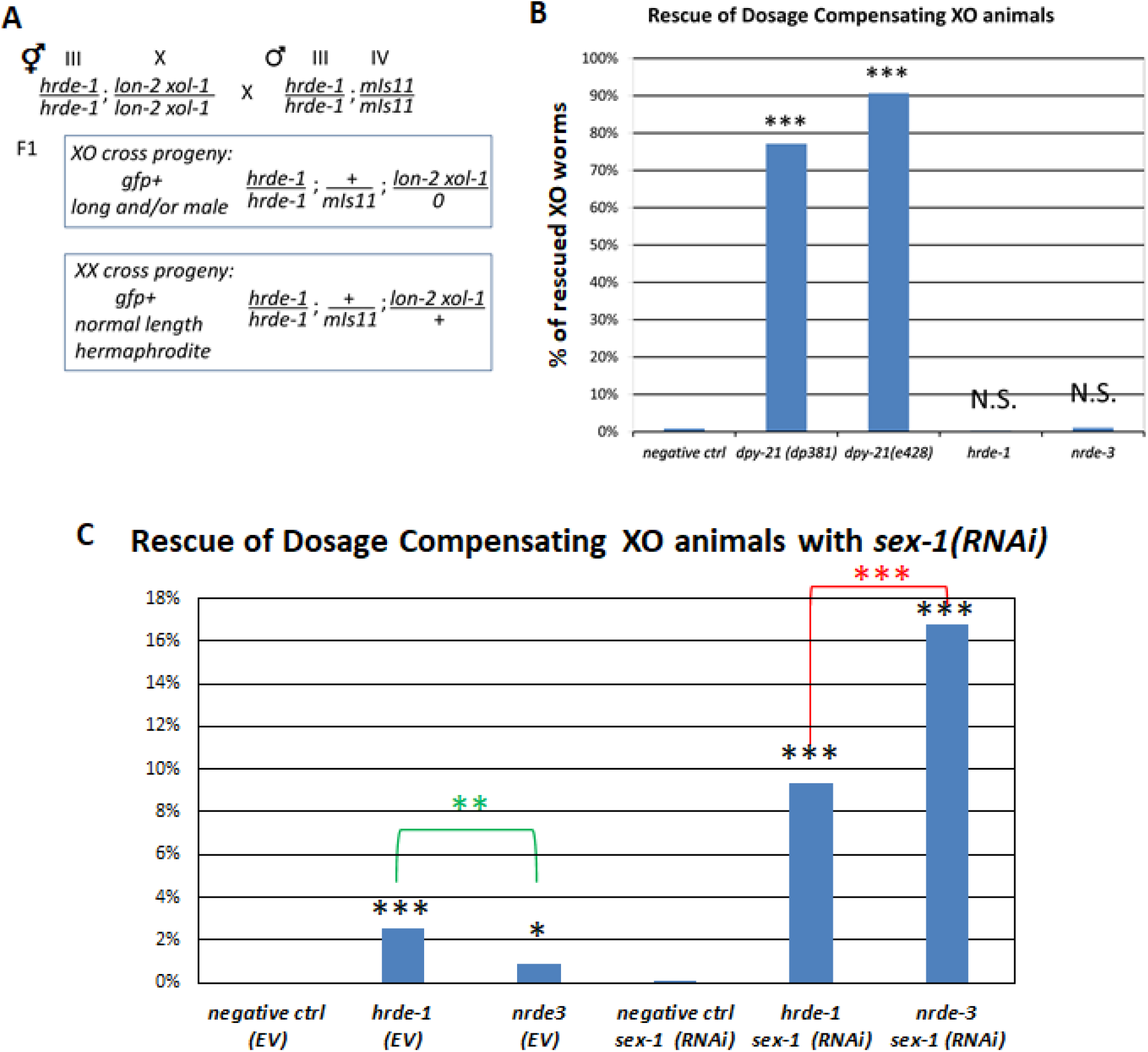
*sex-1* dependent rescue of nuclear Argonaute mutants. **(A)** XO rescue mating cross schematic. *hrde-1* is shown as an example to highlight the experimental condition cross. *hrde-1 Ion-2 xol-1* hermaphrodites are crossed to *hrde-1 mlsll* (GFP+) males. In the F1, GFP+ (*mls11)* worms designate cross progeny. Lon worms represent XO animals hemizygous for Ion-2, some of which develop as male, some as hermaphrodite. Normal length, GFP+ hermaphrodites represent XX cross progeny. **(B)** Percentage of XO worms rescued by various mutations are shown. *dpy-21(e428)* is a null mutation and *dpy-21(dp381)* is a partial loss of function mutation. Both positive controls rescue a large amount of XO animals and *hrde-1* and *nrde-3* on their own do not significantly rescue XO animals. Significance is based on comparison to negative control for each condition. **(C)** Percentage of XO worms rescued by *hrde-1* and *nrde-3* are shown with empty vector or *sex-1(RNAi)* treatment. *hrde-1 sex-l(RNAi)* and *nrde-3 sex-l(RNAi)* rescue a significant portion of XO animals. Black asterisks denote significance for comparison of each condition to negative control of the same RNAi treatment. Significance B and C are based on Chi Square analysis (See Supplemental Figure 2A/B). ***= p<0.001.

### ***nrde-3*** and ***hrde-1*** modulate the structure of a heterochromatin array in somatic cells

It is interesting to note that mutations in *prg-1*, *nrde-3* and *hrde-1* led to decondensation of the X chromosomes, but not chromosome I (Figure 1). This observation suggests that the structure of the X chromosome is particularly sensitive to loss of nuclear RNAi function. To investigate whether non-X chromosome sequences can be affected, we sought to determine whether *nrde-3* and *hrde-1* have a structural impact on a highly heterochromatinized transgenic array (Figure 7). We conducted immunofluorescence staining for the H3K9me3 heterochromatin mark in intestinal cells of *hrde-1* and *nrde-3* mutants and wild type worms bearing the *mIs11* array. Note that this repetitive array is completely silent in intestinal cells. We noticed that the array formed spherical, tightly condensed structures marked by H3K9me3 accumulation in otherwise wild type worms. However, with a *nrde-3* or *hrde-1* mutation in the background, the array appeared less condensed. We characterized these structural defects by binning the heterochromatin array morphology phenotypes into one of four categories. The categories were (in order from most to least condensed): spherical, sickle/bar, starburst, spotted. In wild type, the array tended to take on one of the more compact appearances of spherical (50%) or sickle/bar (16%) H3K9me3 distribution a total of 66% frequency (Figure 7 A and B). In *hrde-1* mutants, the array takes on the spherical (15%) or sickle/bar (25%) H3K9me3 distribution, a total of 40% frequency, and in *nrde-3* mutants the *mIs11* array takes on the spherical (35%) or sickle/bar (18%) H3K9me3 distribution a total of 53% frequency (Figure 7 A and B). Among the two categories of qualitatively decondensed array, the biggest difference for both *hrde-1* and *nrde-3* mutants compared to wild type was a more than two-fold increase of the frequency of starburst H3K9me3 distribution. While the chromosome I controls from our FISH experiments did not capture differences in nuclear volume in the nuclear RNAi AGOs, the H3K9me3 IF data indicate that on a highly heterochromatinized array, a pronounced effect on chromatin structure morphology can be observed, consistent with a previous report indicating a role for nuclear AGOs in compacting chromatin silenced by RNAi (Fields and Kennedy 2019).

**Figure 7.**
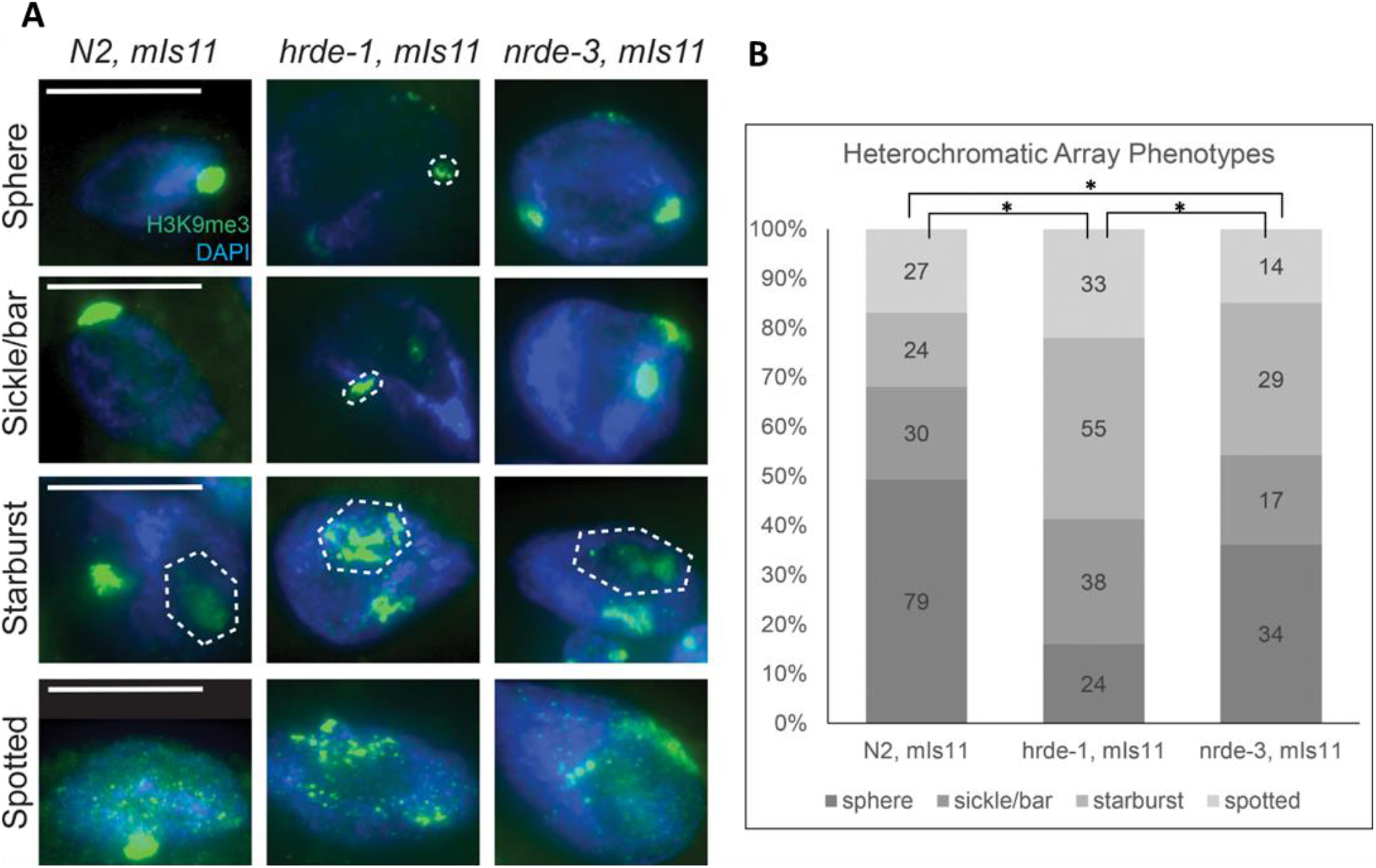
*nrde-3* and *hrde-1* modulate the structure of a heterchromatin array. **(A)** H3K9me3 signal (green) and DAPI (blue) staining is shown in adult hermaphrodite intenstinal cells. Vertical (left) labels represent four qualitative patterns for observed H3K9me3 signal. All patterns were observed for each strain. **(B)** Quantification of qualitative H3K9me3 signal binned by pattern type. Number of nuclei for each pattern are shown within the bars and y-axis shows the proportion of each H3K9me3 pattern as a percentage of that observed in each genotype. *hrde-1* and *nrde-3* exhibited fewer nuclei with the compacted spherical/sickle/bar arrays and a higher proportion of the starburst/spotted arrays. Significance is based on Chi Square analysis for each comparison between genotypes. *= *P*<0.05.

## DISCUSSION

We demonstrated that nuclear RNAi AGOs HRDE-1 and NRDE-3 play dual roles promoting dosage compensation. First, together with the piRNA AGO PRG-1, they repress the master sex determination and dosage compensation switch gene *xol-1*. XOL-1 promotes male development and inhibits dosage compensation, and therefore XOL-1 function must be turned off in hermaphrodites. In addition, HRDE-1, NRDE-3 and PRG-1 also promote dosage compensation downstream of *xol-1*. This downstream role is required for full compaction of dosage compensated X chromosomes, and for healthy development (Figure 8).

**Figure 8.**
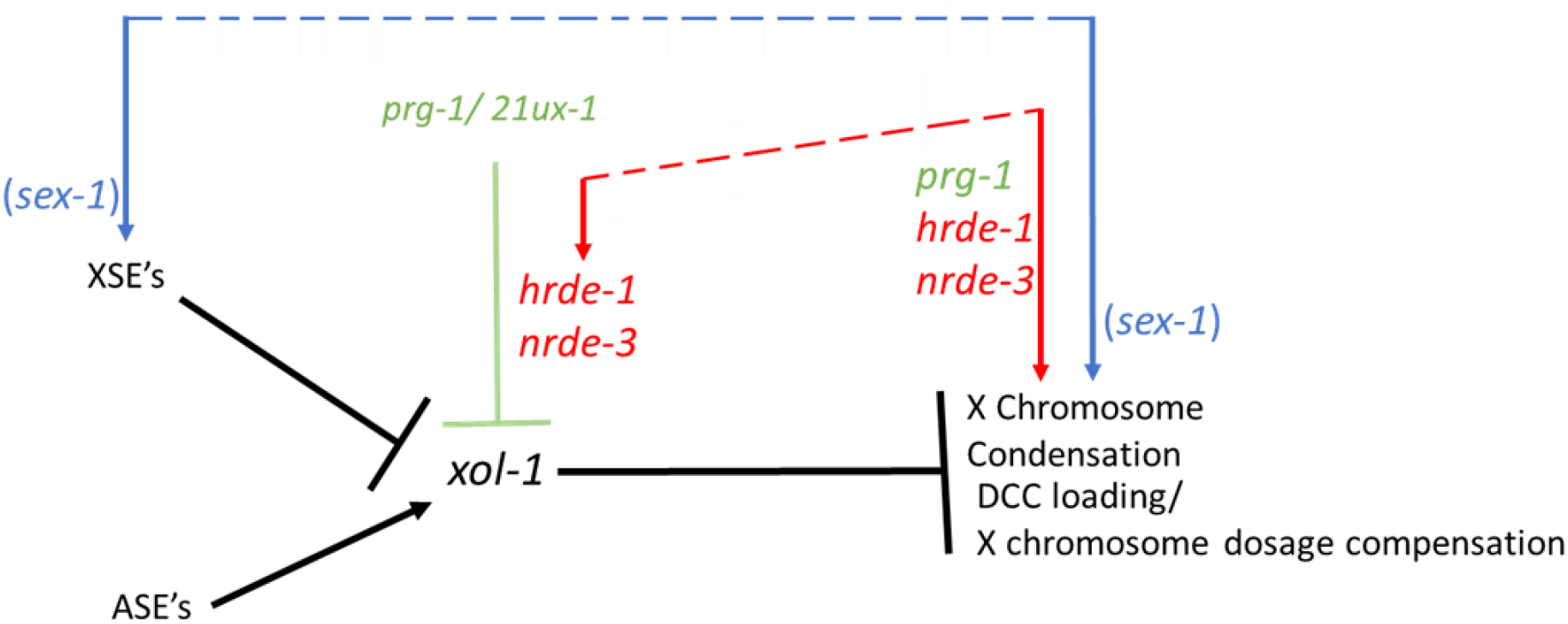
dual role for *hrde-1* and *nrde-3* in dosage compensation. *prg-1* and the *21ux-l* piRNA aid in the repression of *xol-1* in hermaphrodites. *nrde-3* and *hrde-1* also contribute to *xol-1* repression, and cooperate with *sex-1* to contribute to hermaphrodite viability upstream of *xol-1. hrde-1* and *nrde-3* also promote the compaction of X chromosomes, and cooperate with *sex-1* to promote physiological dosage compensation downstream of *xol-1*.

The dual roles of *nrde-3* and *hrde-1* in dosage compensation are similar to the dual roles *sex-1* in their *xol-1*-dependent and independent nature (Carmi *et al.,* 1998, Gladden *et al.,* 2007). While *sex-1’s* role as an X-signal element is well characterized, the *xol-1-*independent role of *sex-1* has not yet been assessed. Identifying genes targeted by *sex-1* will help elucidate how this nuclear hormone receptor homolog gene is acting in conjunction with the nuclear RNAi pathway to promote a significant degree of dosage compensation function. One possibility is that *sex-1*’s role as a nuclear hormone receptor involves recruitment of the nuclear RNAi pathway as one of the mechanism for the repression of its targets. Another possibility is that nuclear RNAi targets a different set of genes that promote dosage compensation in a *sex-1* independent manner. A third possibility is that nuclear RNAi’s influence on chromatin structure augments *sex-1*’s role in promoting dosage compensation. Mutants for the pathway tethering H3K9me3 heterochromatin to the nuclear lamina also exhibit X chromosome decondensation and contribute to dosage compensation in conjunction with *sex-1.* Thus, X chromosome decondensation simultaneously with the loss of *sex-1*’s target gene repression may be what underlies the additive detriment to dosage compensation in double mutants for both *sex-1* and the nuclear RNAi pathway and *sex-1* and the heterochromatin tethering pathway.

While both AGO mutants and *prg-1* exhibit a *xol-1-*independent X chromosome decondensation phenotype to similar degree (Figure 4 A and B), their impacts on male rescue (Figure 2 A), *xol-1* de-repression (Figure 3), hermaphrodite viability (Figure 5) and rescue of XO animals (Figure 6) vary. One measure of the upstream, *xol-1*-dependent role is hermaphrodite viability after *sex-1(RNAi)* treatment. The *xol-1-*dependent hermaphrodite viability of *hrde-1* mutants treated with *sex-1(RNAi)* was only 3%, compared to 16% in *nrde-3 sex-1(RNAi)* (Figure 5 B), indicating that *hrde-1* may play a more prominent role. In contrast, while the degree of *xol-1* de-repression is significantly stronger in both AGO mutants compared to wild type (N2), the level of de-repression is far greater in *nrde-*3 than *hrde-1* (Figure 3). Although impaired hermaphrodite viability is not a direct measure of dosage compensation defect, the synergistic lethality of *hrde-1 sex-1(RNAi)* is *xol-1-*dependent, suggesting that *xol-1* regulation is a major component of *hrde-1*’s function.

Rescue of *xol-1* mutant males by RNAi, rescue of *xol-1* XO animals by introducing genetic mutations, and X chromosome decompaction in *xol-1* mutants, are all measures of the downstream, *xol-1*-independent role. The *xol-1*-independent X chromosome decompaction phenotype was comparable in *prg-1*, *nrde-3* and *hrde-1* mutants (Figure 4). However, the level of X decompaction does not correlate with the level of X derepression, therefore this result should not be interpreted as all genes playing comparable roles (Snyder *et al.,* 2016). The XO rescue experiments on *sex-1(RNAi)* indicate almost a two-fold greater *xol-1-*independent contribution to dosage compensation for *nrde-3* over *hrde-1* (Figure 6 C). However, the degree of *nrde-3* male rescue from the RNAi-based assay in Figure 2 is smaller than that of *hrde-1*. One potential explanation for the discrepancy is that comparisons made between different RNAi conditions are subject to varying degrees of knockdown efficacy between treatments. It is worth noting that the *sex-1(RNAi)* appears to be near (if not) a complete depletion, as the hermaphrodite viability of wild type N2 worms on *sex-1(RNAi)* is close to the published data for *sex-1(null)* mutants (Figure 5 B) (Gladden *et al.,* 2007). *sex-1(RNAi)* is also effective in *hrde-1* and *nrde-3* mutants as well, as evidenced by the severe effects on hermaphrodite viability (Figure 5). Thus, the XO rescue experiments, using loss of function mutants for each AGO and identical RNAi conditions afford a potentially more accurate comparison of contributions from each individual AGO to dosage compensation. In this vein, a bigger role for *nrde-3* downstream of *xol-1* is consistent with its expression pattern in hermaphrodite somatic cells from the embryo through adulthood (Guang *et al.,* 2008). After DCC loading to hermaphrodite X chromosomes (~30-50 cell embryo) NRDE-3’s presence in the soma may reinforce maintenance of dosage compensation through the directing of chromatin modifications (Chuang *et al.,* 1994, Dawes *et al.,* 1999). Future studies utilizing a conditional *nrde-3* depletion to determine the time in development when *nrde-3* is required for dosage compensation will aid in understanding whether *nrde-3* is maintaining and/or initiating the repressive chromatin landscape for dosage compensation.

Considering the germline-enriched expression patterns of *prg-1* and *hrde-1,* the somatic X chromosome decondensation phenotype in both mutant adults is intriguing. The fact that *prg-1* (Tang *et al.,* 2018) and *hrde-1* (Figure 3) repress *xol-1* and independently maintain X chromosome compaction (Figure 4) begs the question of when in development and how mechanistically this contribution is made. Since neither *dcr-1(mg375)* nor *ergo-1(tm1860)* mutants with impaired endogenous primary 26G siRNA accumulation exhibit X chromosome decondensation (Figure 1A and B), the contribution of nuclear AGOs to X chromosome compaction could originate from the piRNA pathway which bypasses the requirement for 26G primary siRNAs but utilizes the nuclear RNAi machinery for silencing (Ashe *et al.,* 2012; Lee *et al.,* 2012; Luteijn *et al.,* 2012; Shirayama *et al.,* 2012). If that were the case, *prg-1* targets may include additional genes important for maintaining the compaction of dosage compensating X chromosomes. The fact that the somatic nuclear AGO *nrde-3* also plays a role both in *xol-1* repression and in *xol-1*-independent X decompaction raises the possibility that repression of some of these *prg-1* targets are maintained by NRDE-3 in the embryo, rather than HRDE-1.

Alternatively, the capacity for these nematode-specific AGOs to direct chromatin modifications may be directly featured in X chromosome compaction. Given *hrde-1’s* chromatin compaction role in the germline and soma (Fields and Kennedy, 2019), its plausible that in *sex-1* mutant hermaphrodites where *xol-1* is de-repressed and dosage compensation is impaired, the loss of *hrde-1*-mediated heterochromatin signatures on the X chromosome further exasperates DCC initiation and/or maintenance. HRDE-1 may reinforce chromatin compaction and direct some degree of heterochromatin formation in the early embryo as well, and this role may then be taken over by NRDE-3 in later stages of embryonic development. The recent discovery of MTR-4’s role in the NRDE complex suggests that there may be additional important genes interacting with the nuclear RNAi machinery (Wan *et al.,* 2020). The NRDE complex is the link between nuclear RNAi target genes, histone modifications, and co-transcriptional silencing (Guang *et al.,* 2008, Guang *et al.,* 2010, Buckley *et al.,* 2011, Ashe *et al.,* 2012, Buckley *et al.,* 2012, Luteijn *et al.,* 2012). Thus, identifying the landscape of NRDE complex binding on *C. elegans* chromosomes and interactions with dosage compensation regulators will further test whether a dosage compensation role is attributed to nuclear RNAi-mediated regulation of genes functioning in dosage compensation, or to nuclear RNAi-mediated regulation of chromosome architecture.

## ACKNOWLEDGEMENTS

This work was supported by the National Institute of General Medical Sciences grant NIH R01GM13385801 to G.C. M.B.D. was partially supported by Michigan Predoctoral Training in Genetics (T32 GM007544), and by the Edwards Fellowship and Okkleberg Fellowship from the Department of MCDB at the University of Michigan. Some strains were provided by the Caenorhabditis Genetics Center (CGC), which is funded by NIH Office of Research Infrastructure Programs (P40 OD010440). We thank Joshua Bembenek for the critical reading and feedback on this manuscript. We also thank all members of the Csankovszki lab for helpful discussions.

## Notes

### Competing Interest Statement

The authors have declared no competing interest.

## REFERENCES

Almeida M. V., M. A. Andrade-Navarro, and R. F. Ketting, 2019 Function and evolution of nematode RNAi pathways. Non-coding RNA 5: 8.

Ambros V., R. C. Lee, A. Lavanway, P. T. Williams, and D. Jewell, 2003 MicroRNAs and other tiny endogenous RNAs in *C. elegans*. Current Biology 13: 807–818.

Albritton S. E., and S. Ercan, 2018 *Caenorhabditis elegans* Dosage Compensation: Insights into Condensin-Mediated Gene Regulation. Trends in Genetics 34: 41–53.

Batista P. J., J. G. Ruby, J. M. Claycomb, R. Chiang, N. Fahlgren, et al., 2008 PRG-1 and 21U-RNAs interact to form the piRNA complex required for fertility in *C. elegans*. Molecular cell 31: 67–78.

Bernstein E., A. A. Caudy, S. M. Hammond, and G. J. Hannon, 2001 Role for a bidentate ribonuclease in the initiation step of RNA interference. Nature 409: 363–366.

Billi A. C., M. A. Freeberg, A. M. Day, S. Y. Chun, V. Khivansara, et al., 2013 A conserved upstream motif orchestrates autonomous, germline-enriched expression of *Caenorhabditis elegans* piRNAs. PLoS genetics 9: e1003392.

Brejc K., Q. Bian, S. Uzawa, B. S. Wheeler, E. C. Anderson, et al., 2017 Dynamic control of X chromosome conformation and repression by a histone H4K20 demethylase. Cell 171: 85–102.e23.

Brenner S., 1974 The genetics of *Caenorhabditis elegans* Genetics 77: 71.

Broverman S. A., and P. M. Meneely, 1994 Meiotic mutants that cause a polar decrease in recombination on the X chromosome in *Caenorhabditis elegans*. Genetics 136: 119–127.

Buckley B. A., K. B. Burkhart, S. G. Gu, G. Spracklin, A. Kershner, et al., 2012 A nuclear Argonaute promotes multigenerational epigenetic inheritance and germline immortality. Nature 489: 447–451.

Burkhart K. B., S. Guang, B. A. Buckley, L. Wong, A. F. Bochner, et al., 2011 A Pre-mRNA– associating factor links endogenous siRNAs to chromatin regulation. PLoS genetics 7: e1002249.

Carmi I., J. B. Kopczynski, and B. J. Meyer, 1998 The nuclear hormone receptor SEX-1 is an X-chromosome signal that determines nematode sex. Nature 396: 168–173.

Chuang P.-T., D. G. Albertson, and B. J. Meyer, 1994 DPY-27: a chromosome condensation protein homolog that regulates *C. elegans* dosage compensation through association with the X chromosome. Cell 79: 459–474.

Conine C. C., P. J. Batista, W. Gu, J. M. Claycomb, D. A. Chaves, et al., 2010 Argonautes ALG-3 and ALG-4 are required for spermatogenesis-specific 26G-RNAs and thermotolerant sperm in *Caenorhabditis elegans*. Proceedings of the National Academy of Sciences 107: 3588–3593.

Consortium C. elegans D. M., 2012 Large-scale screening for targeted knockouts in the *Caenorhabditis elegans* genome. G3: Genes Genomes Genetics 2: 1415–1425.

Crane E., Q. Bian, R. P. McCord, B. R. Lajoie, B. S. Wheeler, et al., 2015 Condensin-driven remodelling of X chromosome topology during dosage compensation. Nature 523: 240–244.

Csankovszki G., P. McDonel, and B. J. Meyer, 2004 Recruitment and spreading of the *C. elegans* dosage compensation complex along X chromosomes. Science 303: 1182–1185.

Csankovszki G., K. Collette, K. Spahl, J. Carey, M. Snyder, et al., 2009 Three distinct condensin complexes control *C. elegans* chromosome dynamics. Current Biology 19: 9–19.

Das P. P., M. P. Bagijn, L. D. Goldstein, J. R. Woolford, N. J. Lehrbach, et al., 2008 Piwi and piRNAs act upstream of an endogenous siRNA pathway to suppress Tc3 transposon mobility in the *Caenorhabditis elegans* germline. Molecular cell 31: 79–90.

Dawes H. E., D. S. Berlin, D. M. Lapidus, C. Nusbaum, T. L. Davis, et al., 1999 Dosage Compensation Proteins Targeted to X Chromosomes by a Determinant of Hermaphrodite Fate. Science 284: 1800.

Fields B. D., and S. Kennedy, 2019 Chromatin compaction by small RNAs and the nuclear RNAi machinery in *C. elegans*. Scientific reports 9: 1–9.

Gent J. I., M. Schvarzstein, A. M. Villeneuve, S. G. Gu, V. Jantsch, et al., 2009 A *Caenorhabditis elegans* RNA-directed RNA polymerase in sperm development and endogenous RNA interference. Genetics 183: 1297–1314.

Gent J. I., A. T. Lamm, D. M. Pavelec, J. M. Maniar, P. Parameswaran, et al., 2010 Distinct phases of siRNA synthesis in an endogenous RNAi pathway in *C. elegans* soma. Molecular cell 37: 679–689.

Gladden J. M., B. Farboud, and B. J. Meyer, 2007 Revisiting the X:A Signal That Specifies *Caenorhabditis elegans* Sexual Fate. Genetics 177: 1639–1654.

Gu W., M. Shirayama, D. Conte Jr, J. Vasale, P. J. Batista, et al., 2009 Distinct argonaute-mediated 22G-RNA pathways direct genome surveillance in the *C. elegans* germline. Molecular cell 36: 231–244.

Gu S. G., J. Pak, S. Guang, J. M. Maniar, S. Kennedy, et al., 2012 Amplification of siRNA in *Caenorhabditis elegans* generates a transgenerational sequence-targeted histone H3 lysine 9 methylation footprint. Nature genetics 44: 157–164.

Guang S., A. F. Bochner, D. M. Pavelec, K. B. Burkhart, S. Harding, et al., 2008 An Argonaute transports siRNAs from the cytoplasm to the nucleus. Science 321: 537–541.

Guang S., A. F. Bochner, K. B. Burkhart, N. Burton, D. M. Pavelec, et al., 2010 Small regulatory RNAs inhibit RNA polymerase II during the elongation phase of transcription. Nature 465: 1097–1101.

Han T., A. P. Manoharan, T. T. Harkins, P. Bouffard, C. Fitzpatrick, et al., 2009 26G endo-siRNAs regulate spermatogenic and zygotic gene expression in *Caenorhabditis elegans*. Proceedings of the National Academy of Sciences 106: 18674–18679.

Höck J., and G. Meister, 2008 The Argonaute protein family. Genome biology 9: 1–8.

Hoogewijs D., K. Houthoofd, F. Matthijssens, J. Vandesompele, and J. R. Vanfleteren, 2008 Selection and validation of a set of reliable reference genes for quantitative sod gene expression analysis in *C. elegans*. BMC molecular biology 9: 1–8.

Jans J., J. M. Gladden, E. J. Ralston, C. S. Pickle, A. H. Michel, et al., 2009 A condensin-like dosage compensation complex acts at a distance to control expression throughout the genome. Genes & development 23: 602–618.

Kramer M., A.-L. Kranz, A. Su, L. H. Winterkorn, S. E. Albritton, et al., 2016 Correction: developmental dynamics of X-chromosome dosage compensation by the DCC and H4K20me1 in *C. elegans*. PLoS genetics 12: e1005899.

Kruesi W. S., L. J. Core, C. T. Waters, J. T. Lis, and B. J. Meyer, 2013 Condensin controls recruitment of RNA polymerase II to achieve nematode X-chromosome dosage compensation, (N. Proudfoot, Ed.). eLife 2: e00808.

Lau A. C., K. Nabeshima, and G. Csankovszki, 2014 The *C. elegans* dosage compensation complex mediates interphase X chromosome compaction. Epigenetics & chromatin 7: 1–16.

Lau A. C., and G. Csankovszki, 2015 Balancing up and downregulation of the *C. elegans* X chromosomes. Current opinion in genetics & development 31: 50–56.

Lee H.-C., W. Gu, M. Shirayama, E. Youngman, D. Conte Jr, et al., 2012 *C. elegans* piRNAs mediate the genome-wide surveillance of germline transcripts. Cell 150: 78–87.

Luteijn M. J., P. van Bergeijk, L. J. T. Kaaij, M. V. Almeida, E. F. Roovers, et al., 2012 Extremely stable Piwi-induced gene silencing in *Caenorhabditis elegans*. EMBO J 31: 3422–3430.

Mao H., C. Zhu, D. Zong, C. Weng, X. Yang, et al., 2015 The Nrde pathway mediates small-RNA-directed histone H3 lysine 27 trimethylation in *Caenorhabditis elegans*. Current Biology 25: 2398–2403.

Miller L. M., J. D. Plenefisch, L. P. Casson, and B. J. Meyer, 1988 xol-1: a gene that controls the male modes of both sex determination and X chromosome dosage compensation in *C. elegans*. Cell 55: 167–183.

Nabeshima K., S. Mlynarczyk-Evans, and A. M. Villeneuve, 2011 Chromosome painting reveals asynaptic full alignment of homologs and HIM-8–dependent remodeling of X chromosome territories during *Caenorhabditis elegans* meiosis. PLoS genetics 7: e1002231.

Ohno S., 1967 Sex chromosome and sex-linked genes Springer. Chromosoma 23: 1–9.

Pak J., and A. Fire, 2007 Distinct Populations of Primary and Secondary Effectors During RNAi in *C. elegans*. Science 315: 241–244.

Parhad S. S., and W. E. Theurkauf, 2019 Rapid evolution and conserved function of the piRNA pathway. Royal Society Open Biology 9: 180181.

Pavelec D. M., J. Lachowiec, T. F. Duchaine, H. E. Smith, and S. Kennedy, 2009 Requirement for the ERI/DICER complex in endogenous RNA interference and sperm development in *Caenorhabditis elegans*. Genetics 183: 1283–1295.

Petty E., E. Laughlin, and G. Csankovszki, 2011 Regulation of DCC localization by HTZ-1/H2A. Z and DPY-30 does not correlate with H3K4 methylation levels. PLoS One 6: e25973.

Pferdehirt R. R., W. S. Kruesi, and B. J. Meyer, 2011 An MLL/COMPASS subunit functions in the *C. elegans* dosage compensation complex to target X chromosomes for transcriptional regulation of gene expression. Genes & Development 25: 499–515.

Phillips C. M., C. Wong, N. Bhalla, P. M. Carlton, P. Weiser, et al., 2005 HIM-8 binds to the X chromosome pairing center and mediates chromosome-specific meiotic synapsis. Cell 123: 1051–1063.

Powell J. R., M. M. Jow, and B. J. Meyer, 2005 The T-Box Transcription Factor SEA-1 Is an Autosomal Element of the X:A Signal that Determines *C. elegans* Sex. Developmental Cell 9: 339–349.

Reed K. J., J. M. Svendsen, K. C. Brown, B. E. Montgomery, T. N. Marks, et al., 2020 Widespread roles for piRNAs and WAGO-class siRNAs in shaping the germline transcriptome of *Caenorhabditis elegans*. Nucleic acids research 48: 1811–1827.

Reinke V., I. S. Gil, S. Ward, and K. Kazmer, 2004 Genome-wide germline-enriched and sex-biased expression profiles in *Caenorhabditis elegans*. Development 131(2): 311–323.

Rhind N. R., L. M. Miller, J. B. Kopczynski, and B. J. Meyer, 1995 xo1-1 acts as an early switch in the *C. elegans* male/hermaphrodite decision. Cell 80: 71–82.

Rice W. R., 1984 Sex chromosomes and the evolution of sexual dimorphism. Evolution 735–742.

Ruby J. G., C. Jan, C. Player, M. J. Axtell, W. Lee, et al., 2006 Large-scale sequencing reveals 21U-RNAs and additional microRNAs and endogenous siRNAs in *C. elegans*. Cell 127: 1193–1207.

Simon M., P. Sarkies, K. Ikegami, A.-L. Doebley, L. D. Goldstein, et al., 2014 Reduced insulin/IGF-1 signaling restores germ cell immortality to *Caenorhabditis elegans* Piwi mutants. Cell reports 7: 762–773.

Snyder M. J., A. C. Lau, E. A. Brouhard, M. B. Davis, J. Jiang, et al., 2016 Anchoring of heterochromatin to the nuclear lamina reinforces dosage compensation-mediated gene repression. PLoS genetics 12: e1006341.

Strome S., W. G. Kelly, S. Ercan, and J. D. Lieb, 2014 Regulation of the X Chromosomes in *Caenorhabditis elegans*. Cold Spring Harb Perspect Biol 6: a018366.

Tabara H., M. Sarkissian, W. G. Kelly, J. Fleenor, A. Grishok, et al., 1999 The rde-1 gene, RNA interference, and transposon silencing in *C. elegans*. Cell 99: 123–132.

Tang W., M. Seth, S. Tu, E.-Z. Shen, Q. Li, et al., 2018 A sex chromosome piRNA promotes robust dosage compensation and sex determination in *C. elegans*. Developmental cell 44: 762–770. e3.

Torres E. M., B. R. Williams, and A. Amon, 2008 Aneuploidy: cells losing their balance. Genetics 179: 737–746.

Vasale J. J., W. Gu, C. Thivierge, P. J. Batista, J. M. Claycomb, et al., 2010 Sequential rounds of RNA-dependent RNA transcription drive endogenous small-RNA biogenesis in the ERGO-1/Argonaute pathway. Proceedings of the National Academy of Sciences 107: 3582–3587.

Vielle A., J. Lang, Y. Dong, S. Ercan, C. Kotwaliwale, et al., 2012 H4K20me1 Contributes to Downregulation of X-Linked Genes for *C. elegans* Dosage Compensation. PLOS Genetics 8: e1002933.

Wan G., J. Yan, Y. Fei, D. J. Pagano, and S. Kennedy, 2020 A Conserved NRDE-2/MTR-4 Complex Mediates Nuclear RNAi in *Caenorhabditis elegans*. Genetics 216: 1071–1085.

Weiser N. E., D. X. Yang, S. Feng, N. Kalinava, K. C. Brown, et al., 2017 MORC-1 integrates nuclear RNAi and transgenerational chromatin architecture to promote germline immortality. Developmental cell 41: 408–423.

Welker N. C., D. M. Pavelec, D. A. Nix, T. F. Duchaine, S. Kennedy, et al., 2010 Dicer’s helicase domain is required for accumulation of some, but not all, *C. elegans* endogenous siRNAs. Rna 16: 893–903.

Wells M. B., M. J. Snyder, L. M. Custer, and G. Csankovszki, 2012 *Caenorhabditis elegans* dosage compensation regulates histone H4 chromatin state on X chromosomes. Molecular and cellular biology 32: 1710–1719.

Yigit E., P. J. Batista, Y. Bei, K. M. Pang, C.-C. G. Chen, et al., 2006 Analysis of the *C. elegans* Argonaute family reveals that distinct Argonautes act sequentially during RNAi. Cell 127: 747–757.

