## Supplementary figures and images for "Dual Roles for Nuclear RNAi Argonautes in *C. elegans* Dosage Compensation"

### Supplemental Tables 1 and 2

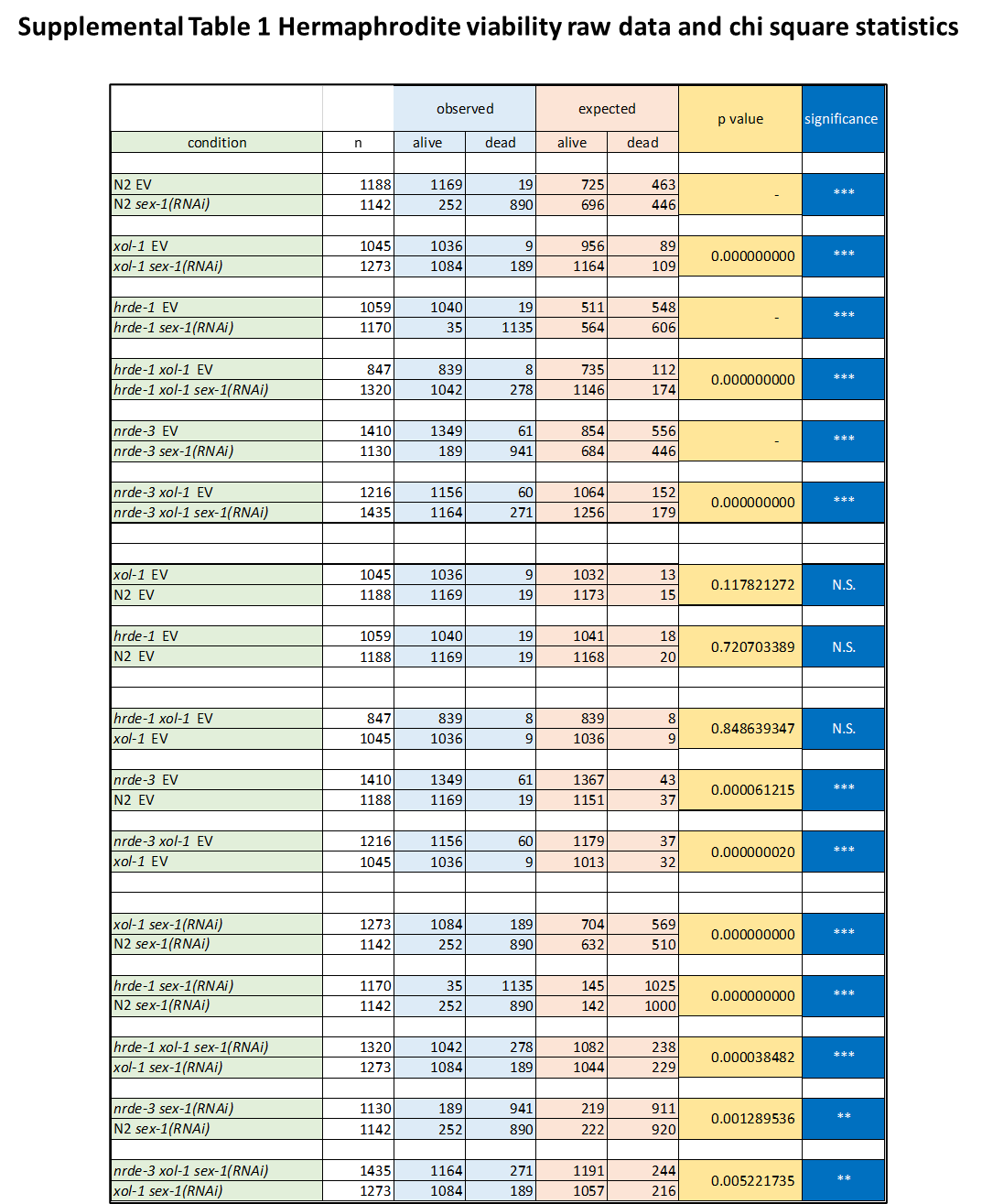

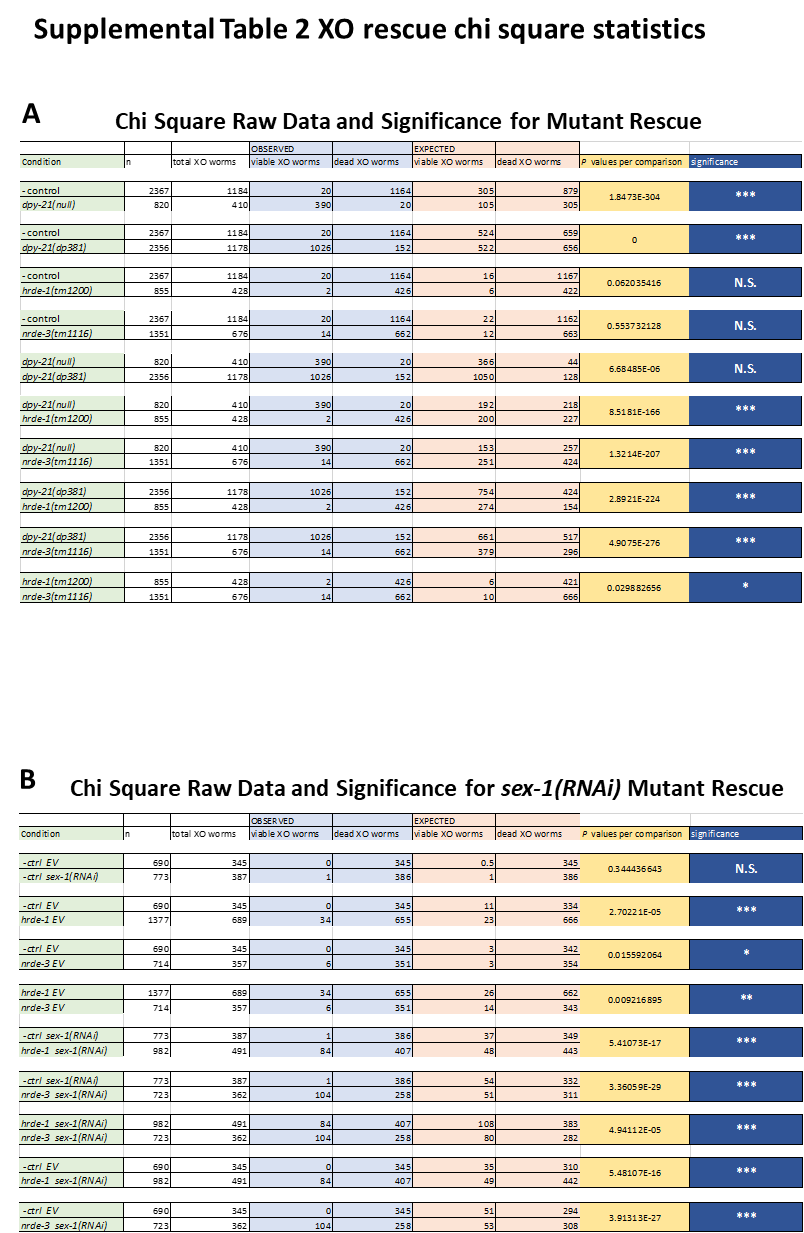
